# NCBI BLAST + integrated into Galaxy

**DOI:** 10.1101/014043

**Authors:** Peter J. A. Cock, John M. Chilton, Björn Grüning, James E. Johnson, Nicola Soranzo

## Abstract

**Background:** The NCBI BLAST suite has become ubiquitous in modern molecular biology, used for small tasks like checking capillary sequencing results of single PCR products through to genome annotation or even larger scale pan-genome analyses. For early adopters of the Galaxy web-based biomedical data analysis platform, integrating BLAST was a natural step for sequence comparison workflows.

**Findings:** The command line NCBI BLAST+ tool suite was wrapped for use within Galaxy, defining appropriate datatypes as needed, with the goal of making common BLAST tasks easy, and advanced tasks possible.

**Conclusions:** This effort has been come an informal international collaborative effort, and is deployed and used on Galaxy servers worldwide. Several example use-cases are described herein.

## Findings

### Introduction

The Basic Local Alignment Search Tool (BLAST) [1] has arguably become the best known and most widely used bioinformatics tool in molecular biology. Indeed, BLAST is now so ubiquitous that, like PCR (polymerase chain reaction), it has become both a noun and verb in the patois of molecular biology, with the acronym rarely spelt out, and unfortunately frequently used without citation.

In our opinions, a key factor in the widespread adoption of BLAST has been the easy to use NCBI hosted BLAST web-server, providing (sufficiently) quick search results against regularly updated global sequence databases. The NCBI BLAST web interface is designed for performing one query at a time, meaning larger searches have to be automated for batch processing within a script or by running BLAST as a command line program. Moreover, with larger BLAST output datasets, automation also became increasingly important for the analysis of BLAST output. These needs led to community developed libraries like BioPerl [2], Biopython [3], BioJava [4] and BioRuby [5] including code for calling BLAST and parsing its output. While greatly facilitating scripted BLAST workflows, large scale BLAST analysis still required a broad bioinformatics skill set including programming, dealing with complex file types, and working at the command line.

With the advent of “next generation” high-throughput sequencing technology, the falling cost of sequence data generation has resulted in a data abundance and all-too-often analysis bottlenecks. This life science “informatics crisis” was one of the motivations behind the Galaxy Project which provides a platform for running a broad collection of bioinformatics tools via a consistent web-interface [6, 7].

From the Galaxy end user’s perspective, no local software is required other than a recent web-browser, yet they can run multiple bioinformatics tools (which may be Linux specific) from their desktop, and easily chain together the output of one tool as the input of another. Moreover, Galaxy’s workflow feature allows users to create and share repeatable analysis pipelines. To encourage reproducibility these can be published as part of the methods in a scientific paper, or in a repository like myExperiment [8].

Galaxy is an open source project, and an international development community has grown up contributing improvements both to the core software, and more importantly to a growing pool of new tools and datatype definitions which can be added to individual Galaxy servers. These extensions are typically shared via the Galaxy Tool Shed (https://usegalaxy.org/toolshed or https://toolshed.g2.bx.psu.edu), which is a public repository of tools and workflows [9], from where they can then be installed on individual Galaxy servers. A few examples were recently published [10, 11, 12, 13].

The expansion of a Galaxy developer community outside the project core team has been facilitated by much of Galaxy’s development being coordinated online in public, using mailing lists, source code repositories (https://bitbucket.org/galaxy/ hosted by Atlassian, and https://github.com/galaxyproject/ hosted by GitHub, Inc.) and project management tools (tracking issues and feature requests hosted by Trello Inc). Moreover it has been supported by an annual Galaxy Community Conference since 2011, and full time staff on the Galaxy Project dedicated to outreach work helping to nurture an engaged Galaxy user-community.

While a free to use public server is available at https://usegalaxy.org at Pennsylvania State University, many groups and institutes run their own Galaxy servers. This allows customisation of Galaxy with additional tools of local interest, control of potentially sensitive data, and allows exploitation of local computing infrastructure, or even rented computers from a Cloud Computing provider such as Amazon Web Services (AWS) using Galaxy CloudMan [14]. Furthermore public Galaxy servers are now also being provided by groups wishing to make their own tools immediately available to run by the wider community, avoiding the need to write a bespoke web-interface [13, 15, 11].

This manuscript describes our NCBI BLAST+ [16] wrappers for Galaxy, and associated tools and datatype definitions. Currently these tools have not been made available at the https://usegalaxy.org public server due to concerns over the resulting computational load (J. Taylor, personal communication, 2013). However, they are available from the Galaxy Tool Shed for automated installation into a local Galaxy instance, or from our source code repository (https://github.com/peterjc/galaxy_blast/ hosted by GitHub Inc.), and are released under the open source MIT licence.

### Results

Tables 1 and 2 summarise the Galaxy tools described, wrappers for the NCBI BLAST+ command line tools, and BLAST related tools respectively. Table 3 summarises the Galaxy datatypes used or defined. We now describe some use-cases and workflows combining these tools within Galaxy. Further examples were described in [10].

**Table 1.** NCBI BLAST+ Galaxy tools described. Each table row represents a separate Galaxy tool, all available from https://toolshed.g2.bx.psu.edu/view/devteam/ncbi_blast_plus/ on the Galaxy Tool Shed. Typically there is a separate Galaxy Tool for each different underlying NCBI BLAST+ command line tool, however the main functions of the blastdbcmd command line tool are represented as two separate Galaxy tools. We intend to add further wrappers later, including for the command line tools psiblast and deltablast.

| Galaxy Tool Name | Description | Reference(s) |
| --- | --- | --- |
| NCBI BLAST+ blastp | Protein vs. protein | [1, 16] |
| NCBI BLAST+ blastn | Nucleotide vs. nucleotide | [1, 16] |
| NCBI BLAST+ blastx | Translated nucleotide vs. protein | [1, 16] |
| NCBI BLAST+ tblastn | Protein vs. translated nucleotide | [1, 16] |
| NCBI BLAST+ tblastx | Translated nucleotide vs. translated nucleotide | [1, 16] |
| NCBI BLAST+ makeblastdb | Make BLAST nucleotide or protein database | [1, 16] |
| NCBI BLAST+ makeprofiledb | Make BLAST protein domain database | [1, 16] |
| NCBI BLAST+ blastdbcmd entry(s) | Extract sequence(s) from BLAST database | [1, 16] |
| NCBI BLAST+ blastdbcmd info | Show BLAST database information | [1, 16] |
| NCBI BLAST+ dustmasker | Nucleotide masking using DUST algorithm | [1, 16] |
| NCBI BLAST+ segmasker | Protein masking using SEG algorithm | [1, 16] |
| NCBI BLAST+ windowmasker | Window based sequence masker | [1, 16] |
| NCBI BLAST+ convert2blastmask | Lowercase masking | [1, 16] |
| NCBI BLAST+ rpsblast | Protein vs. protein domain | [16, 31] |
| NCBI BLAST+ rpstblastn | Translated nucleotide vs. protein domain | [16, 31] |

**Table 2.** Additional Galaxy tools using NCBI BLAST+. Each table row represents a separate Galaxy tool, all available from the Galaxy Tool Shed (https://toolshed.g2.bx.psu.edu/view/).

| URL, Galaxy Tool Name | Description | Reference(s) |
| --- | --- | --- |
| https://toolshed.g2.bx.psu.edu/view/devteam/ncbi_blast_plus/ |  |  |
| BLAST XML to tabular | Convert BLAST XML output into tabular output | [10] |
| https://toolshed.g2.bx.psu.edu/view/peterjc/blast_rbh/ |  |  |
| BLAST Reciprocal Best Hits (RBH) | Takes two FASTA inputs, returns table |  |

**Table 3.** Galaxy datatypes used or defined. Each table row represents a separate Galaxy datatype, either available from https://toolshed.g2.bx.psu.edu/view/devteam/blast_datatypes/ on the Galaxy Tool Shed, or already built into Galaxy.

| Galaxy Datatype | Type | Description |
| --- | --- | --- |
| tabular | Built-in | Tab-separated plain text table, used as default BLAST+ output |
| text | Built-in | Plain text, used for human-readable BLAST+ output |
| html | Built-in | Webpage, used for human-readable BLAST+ output with hyper-links |
| blastxml | Add-on | BLAST XML output |
| blastdbn | Add-on | BLAST database of nucleotide sequences, e.g. for BLASTN |
| blastdbp | Add-on | BLAST database of protein sequences, e.g. for BLASTP |
| blastdbd | Add-on | BLAST database of protein domain PSSMs, e.g. for RPS-BLAST |
| maskinfo-asn1 | Add-on | BLAST masking information files as text ASN.1 |
| maskinfo-asn1-binary | Add-on | BLAST masking information files as binary ASN.1 |

#### Assessing a de novo assembly

While more specialised tools exist for the annotation of a *de novo* assembly (e.g. Augustus [17], Glimmer3 [18] and Prokka [19], which we previously wrapped for use in Galaxy [10, 13]), BLAST is often used for a first pass assessment. The following example is based on a procedure which a local sequencing service, Edinburgh Genomics, had adopted as part of their quality control (later extended as described in [20]).

- Upload or import Illumina reads in FASTQ format.
- Run a fast assembler like the CLC Assembly Cell (CLC bio, Aarhus, Denmark) which we have wrapped for use within Galaxy (https://toolshed.g2.bx.psu.edu/view/peterjc/clc_assembly_cell) to generate an initial set of contigs.
- Compare these initial contigs to the NCBI NR database using BLASTX, requesting at most one hit and tabular output including the taxonomy fields (and optionally the hit description).

Since the CLC Assembly Cell software is proprietary, our examplar workflow, available from the Galaxy Tool Shed (http://toolshed.g2.bx.psu.edu/view/peterjc/blast_top_hit_species) and myExperiment (http://www.myexperiment.org/workflows/4637.html), starts from a previously generated or imported transcriptome assembly. This analyses a sample of 1000 sequences only, and uses Galaxy data manipulation tools to produce a sorted tally table of species hits suitable for visualisation within Galaxy as a Pie Chart.

This simple taxon assignment can detect obvious contamination, or sample mixup. However, this kind of simple “Top BLAST hit” analysis should be treated with caution due to the potential for spurious matches, or matches to misannotated sequences such as contaminants in published whole genome shotgun assemblies (see for example [21] and reference therein).

#### Finding genes of interest in a de novo assembly

As sequencing costs have fallen, for many organisms is it now practical to sequence the entire genome when interested primarily in a single gene family. In this situation BLAST might be used within Galaxy as follows:

- Upload or import the (meta-)genome or transcriptome assembly in FASTA format.
- Upload protein (or nucleotide) sequence of the gene(s) of interest.
- Run the makeblastdb wrapper to create a BLAST nucleotide database from the assembly.
- Run the blastx (or blastn) wrapper using the gene(s) of interest as the query against the new database.
- Filter the matching contigs from the assembly FASTA using https://toolshed.g2.bx.psu.edu/view/peterjc/seq_filter_by_id (or similar).

If required, rather than extracting complete contigs, Galaxy has tools for working with genomic intervals that could be used to select the matched regions only, as in the next example.

#### Identifying candidate genes clusters

Identification and analysis of gene clusters is an important task in synthetic biology [22, 23]. Unfortunately, identifying candidate gene clusters is quite complex and can take hours for a single genome. However with prior knowledge about the expected genes in a cluster the genome can be screened limiting the search space dramatically.

For this use case a workflow was constructed to query two translated protein sequences against a BLAST nucleotide database for the target genome ([23]; Figure 1). This workflow is available with sample data via the Galaxy Tool Shed (http://toolshed.g2.bx.psu.edu/view/bgruening/find_genes_located_nearby_workflow) and myExperiment (http://www.myexperiment.org/workflows/4584.html).

**Figure 1.**
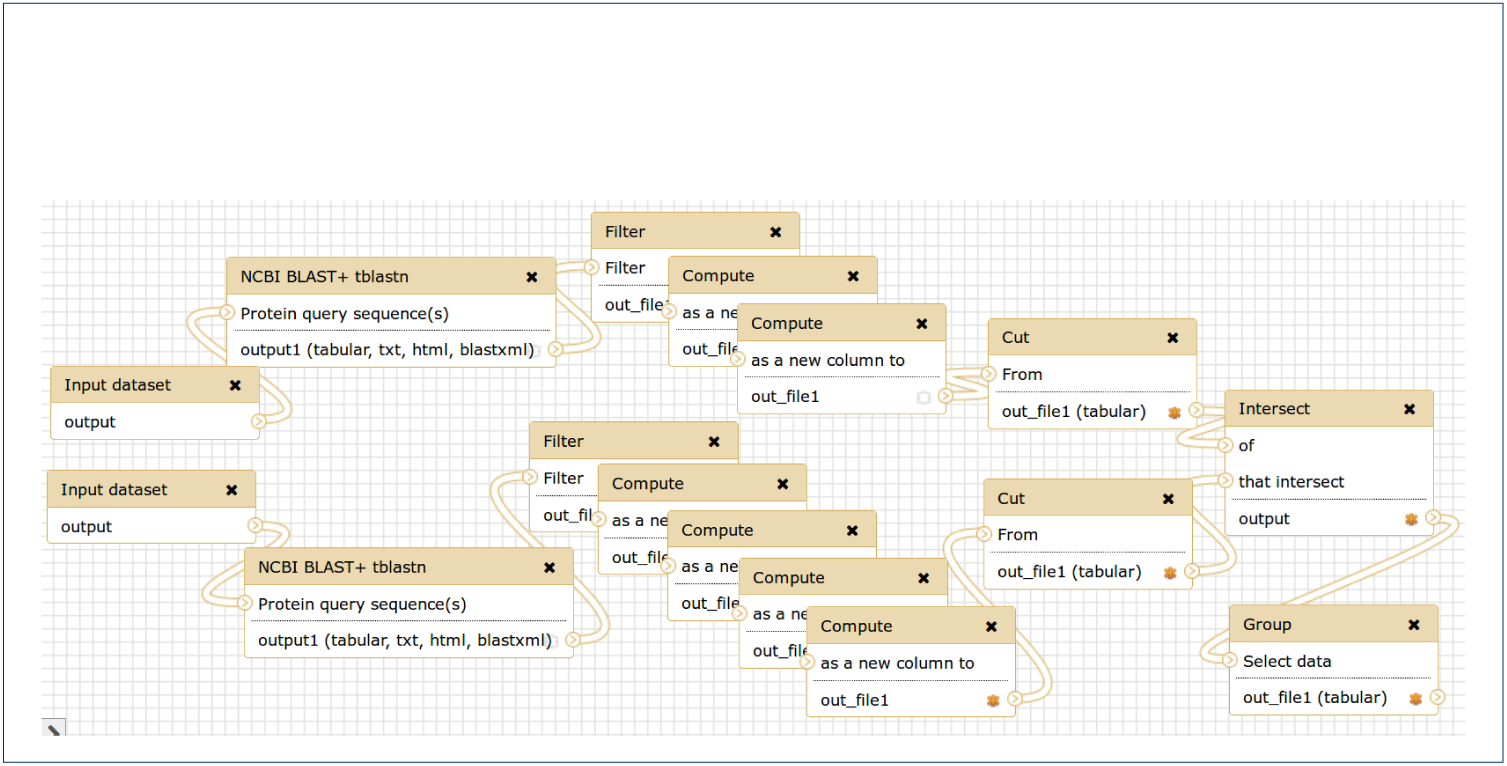
Galaxy Workflow for finding gene clusters. This is a screenshot from the Galaxy Workflow Editor, showing a published example workflow [23] discussed in Results. Given two protein sequences, regions of a genome of interest are identified which contain tblastn matches to both, giving candidate gene clusters for further study.

The TBLASTN results are processed with standard Galaxy text manipulation tools to extract the target sequence identifier and the hits start and stop coordinates. The obtained 3 column interval format is BED-like and the sequence identifier corresponds to the chromosome or contig name. Before intersecting the hit regions we extend one of them by 10, 000 bp up- and downstream, by adding and subtracting 10, 000 from the start and end coordinates, respectively. The intersect tool will work on genomic coordinates, identifying overlapping regions. These regions encode similar proteins to the query sequence, and other proteins in close proximity (< 10, 000 bp). The optional and last step in this example will group and count all sequence identifiers, returning a list of all identified pairs, located nearby and their count.

This approach screens two proteins against all nucleotide sequence from the NCBI nt database within hours on our cluster, leading to all organisms with an interesting gene structure for further investigation. As usual in Galaxy workflows every parameter, including the proximity distance, can be changed and additional steps can be easily added. For example additional filtering to refine the initial BLAST hits, or inclusion of a third query sequence.

#### Identifying novel proteins

Proteogenomics combines genomic information with mass spectrometry derived experimental data for proteomic analysis. To search for evidence of novel proteins, the databases for proteomics search applications are generated from 6-frame translations of genomics or transcripts sequences or cDNA transcripts. With such large databases, proteomics search applications generate a large number of peptide spectral matches (PSMs). The University of Minnesota developed workflows in Galaxy-P (https://usegalaxyp.org/) to automate proteogenomic analysis [24]. These workflows employ the NCBI BLAST+ wrappers to compare the PSM peptides to known proteins in order to filter the PSM list for those more likely to be novel. An additional BLASTP wrapper was deployed in Galaxy-P to use the remote-search option of BLASTP to perform taxon specific searches at NCBI.

### Implementation

Despite its maturity, the Galaxy platform has continued to evolve rapidly, especially in the area of tool definition and distribution. The recently published Galaxy Tool Shed [9] allows anyone hosting a Galaxy instance to install tools and defined dependencies with a few clicks right from the Galaxy web application itself. The NCBI BLAST+ tools described here were among the first tools migrated to the tool shed and have served as drivers of tool shed features and representative examples of how easy it can be to deploy very powerful tools using Galaxy.

The Galaxy BLAST+ wrappers are developed as an open source project using the distributed version control system Git (http://git-scm.com/). We use the hosting service github.com provided by GitHub, Inc., which has become the hub of a growing software development ecosystem. One particular example of this is the continuous integration service travis-ci.org offered by Travis CI GmbH. While complex to setup, every time our source code is updated on GitHub, Travis CI automatically creates a Linux virtual machine, installs BLAST+, the latest Galaxy code, and our wrappers whose functional tests are then run (https://travis-ci.org/peterjc/galaxy_blast). This provides us prompt feedback, where many errors can be caught and dealt with (before releasing via the Galaxy Tool Shed). Furthermore, the BLAST+ wrapper tests have been used by the Galaxy development team when working on the Galaxy test framework.

One of the core concepts in Galaxy is that each dataset has a specified datatype or file format, such as FASTA format sequences, or various FASTQ encodings [25]. Each Galaxy tool normally accepts only specific datatypes as input, and will mark its output files with the appropriate datatype. We defined a set of datatype definitions for BLAST ASN.1 files, BLAST XML and the different BLAST database types (see Table 3). Simple datatypes can be defined by subclassing already existing datatypes. In general additional Python code is required, such as defining a sniff function for auto-detection of the datatype when loading files into Galaxy.

Galaxy also supports simple job-splitting which works at the datatype level with input datatypes (such as FASTA) needing to provide a split method, and output datatypes (such as tabular or BLAST XML) needing to provide a merge method. If this job-splitting is enabled, BLAST searches are automatically parallelised by splitting the FASTA query file into chunks, and then merging the output BLAST results. This is done transparently to the user, and allows genome scale BLAST jobs to be spread across a cluster rather than being processed serially, providing a dramatic speedup.

The Galaxy-P project (Minnesota Supercomputing Institute, University of Minnesota) contributed extensions to Galaxy known as tool macros that make it considerably easier to develop and maintain large suites of Galaxy tools by allowing authors to define high-level abstractions describing any aspect of Galaxy’s XML-based tool description language. These abstractions can be composed together and shared across various tools in a suite. In wrapping NCBI+ BLAST tool suite we have made heavy use of this to avoid the duplication of common parameters, command line arguments, and even help text. In addition to removing hundreds of lines of XML, this helps with consistency and maintenance as many changes need only be made once to the macro definition.

While the Galaxy Tool Shed has greatly simplified installation of additional tools to an existing Galaxy Server, doing this “by hand” remains time consuming and reproducibility suffers. However, this can be scripted, while useful for automated testing (as in our TravisCI setup outlined above), this is vital for large scale deployment. In a similar vein to the Galaxy CloudMan project [14] for automated creation of complete virtual machine images running Galaxy, we have explored using the virtual containers technology from Docker Inc. (https://www.docker.com/) for testing and deployment of a Galaxy server complete with additions like the BLAST+ tools. The Galaxy BLAST Docker Image https://registry.hub.docker.com/u/bgruening/galaxy-blast/ offers a complete Galaxy instance with FTP Server, Job Scheduler and BLAST wrappers [26]. Once docker is installed, the command “docker run -p 8080:80 bgruening/galaxy-blast” will download the image, and start a BLAST enabled Galaxy instance on port 8080. Note that the Docker Image does not currently automate installation of any BLAST databases.

One area which remains a burden for the Galaxy administrator is the provision of local copies of BLAST databases (external to Galaxy), such as in-house unpublished datasets, or the main NCBI BLAST databases from ftp://ftp.ncbi.nlm.nih.gov/blast/db/. The locations of these databases (which may be used outside of Galaxy) are listed in simple tabular configuration files (blastdb*.loc) which store a unique identifier key (which Galaxy records), description (shown to the Galaxy user), and the file path to the database (which can be updated if required, for example due to changes in local storage architecture). In future work we hope to use the Galaxy Data Manager Framework [27] to facilitate this.

### Discussion

Over decades the BLAST suite has grown with improvements like gapped searches [28], and additional functionality such as Position-Specific Iterated BLAST (PSI-BLAST) [28, 29] and protein domain searches with Reverse Position-Specific BLAST (RPS-BLAST) [30]. These Position-Specific Score Matrix (PSSM) based tools under-pin the NCBI Conserved Domain Database (CDD) and the associated web-based Conserved Domain Search service (CD-Search) [30, 31]. More recently, the NCBI BLAST team undertook an ambitious rewrite of the BLAST tool suite, converting the existing “legacy” code base written in the C programming language to use the C++ language instead, dubbed BLAST+ [16].

The expansion of the Galaxy wrappers for BLAST+ has followed a similar course. The initial wrappers focussed on the five core tools (BLASTP, BLASTN, BLASTX, TBLASTN, and TBLASTX), and did not allow the creation of custom BLAST databases. Gradually the scope and contributor base has expanded (Tables 1 and 3), particularly since our publication of genome and protein annotation tools [10], supported by the move to a dedicated source code repository on GitHub. This shift to a distributed international team effort followed discussion both online and in person at the Galaxy Community Conference 2013, and reflects the broad usage of the BLAST+ tools within the Galaxy community.

Future work will include additional wrappers for the remaining or new BLAST+ command line tools, exposing additional command line options via the Galaxy interface, and additional output file formats. Developments within Galaxy will also allow new functionality. For example, we hope to build on the Galaxy Visual Analysis Framework [32] to offer graphical representation of BLAST results within Galaxy, like that offered by the NCBI web-service. Similarly, managing local BLAST databases could be facilitated using the Data Manager Framework [27]

By their nature, the Galaxy *.loc files and associated external datasets (such as NCBI BLAST databases) impose both an administrative overhead, and limitations on reproducibility. One problem is versioning of external datasets requires a copy of each revision be maintained with its own entry in Galaxy’s corresponding *.loc file. In the case of the NCBI BLAST databases, this is hampered by the absence of official versioning. Here a date-stamping approach is possible, for example quarterly snapshots if local storage allows. However, the more practical and likely more common approach here is a single live copy of the NCBI BLAST databases, kept up to date automatically with the NCBI provided Perl scripts or similar. Such setups are often already in place on central computer clusters used for bioinformatics. A second issue with using external datasets in Galaxy is they undermine sharing of workflows between Galaxy servers, since any referenced external datasets must also be synchronised. At a practical level this requires consistent naming schemes. For instance, for current versions of the NCBI BLAST databases we recommend that the Galaxy administrator always use the case-sensitive stem of the file name as the key (e.g. use `nr` in blastdb_p.loc to refer to a current version of the NCBI “non-redundant” BLAST protein database).

Locally, running BLAST+ within Galaxy has been particularly useful for multiquery searches, and searching against unpublished data such as draft genomes since both the local administrator and individual users can create databases. However, the biggest user benefits for data processing come when complete workflows can be completed within Galaxy, as in the examples shown.

## Availability and requirements

- **Project name:** Galaxy wrappers for NCBI BLAST+ & related BLAST tools
- **Project home page:** https://github.com/peterjc/galaxy_blast
- **Operating system(s):** Linux (recommend), Mac
- **Programming language:** Python
- **Other requirements:** Galaxy (and dependencies therein), NCBI BLAST+
- **License:** The MIT License (MIT).
- **Any restrictions to use by non-academics:** No

The Galaxy wrappers are also available from the Galaxy Tool Shed https://toolshed.g2.bx.psu.edu/view/devteam/ncbi_blast_plus for installation to an existing Galaxy server, and as part Docker Image https://registry.hub.docker.com/u/bgruening/galaxy-blast/ providing a Galaxy server with the BLAST+ tools preinstalled.

## Availability of supporting data

The data sets supporting the results of this article are available in the Galaxy BLAST repository, https://github.com/peterjc/galaxy_blast (i.e. sample files used for automated functional testing).

## List of abbreviations used

AWS: Amazon Web Services

BED: Browser Extensible Data (a sequence interval file format);

BLAST: Basic Local Alignment Search Tool;

DDBJ: DNA Data Bank of Japan;

EMBL: European Molecular Biology Laboratory;

INSDC: International Nucleotide Sequence Database Collaboration (NCBI/EMBL/DDBJ);

NCBI: National Center for Biotechnology Information (USA);

PSI-BLAST: Position-Specific Iterated BLAST; PSM: Peptide Spectral Match;

PSSM: Position-Specific Score Matrix;

QBLAST: Queued BLAST (an NCBI web-service);

RPS-BLAST: Reverse Position-Specific BLAST;

XML: Extensible Markup Language.

## Competing interests

The authors declare that they have no competing interests.

## Author’s contributions

All authors have made technical contributions to the tools described, and have read, contributed to, and approved the final manuscript. PJAC initiated this work and continues to coordinate development. The co-authors are listed alphabetically by surname.

## Acknowledgements

We thank NCBI BLAST team for their long running work, the Galaxy Team for their assistance and advice, the Galaxy Community for their feedback and suggestions, and the past, present and future contributors to the Galaxy tools described here. Specifically, this includes: Dannon Baker, Daniel Blankenberg, Edward Kirton, Kanwei Li, and Luobin Yang. The three reviewers are thanked for their constructive feedback, Tom Madden, Gianmauro Cuccuru, and in particular Stian Soiland-Reyes for his attention to detail.

PJAC was funded by the Scottish Government’s Rural and Environment Science and Analytical Services (RESAS) Division. BAG was funded by the DFG CRC 992 Medical Epigenetics. JMC and JEJ were supported in part by NSF grant 1147079.

